# Genotype-by-environment interaction analysis for flowering, maturity time and yield in fonio across traditional and prospective production areas in Northern Benin

**DOI:** 10.64898/2026.05.12.724536

**Authors:** Tania L. I. Akponikpè, Elvire L. Sossa, Idrissou Ahoudou, Abdou R. Ibrahim Bio Yerima, Guillaume L. Amadji, Séverine Piutti, Enoch G. Achigan-Dako

## Abstract

In this study, the critical gap in understanding how fonio responds to contrasting pedoclimatic conditions, both within and outside its traditional production areas was addressed. A multi-environment trial was carried out to identify high-yielding genotypes with either broad stability or specific adaptation, thereby enabling targeted varietal recommendations to support the expansion of fonio cultivation into new areas. Randomized complete block design was used in six environments with eleven genotypes to evaluate flowering and maturity times, and grain yield. The Additive Main effect and Multiplicative Interaction and the Genotype main effect and Genotype × Environment interaction biplots revealed a significant effect of the genotype-by-environment interactions on traits, with genotypes B12 and G31 identified as high-yielding, while genotypes M5 and M14 were revealed as early-flowering and maturing. Genotypes M14 and M15 were adapted to all environments and early maturing. Boukoumbe, known as the fonio production area in Benin, was the most desirable for earliness, while Ina was the most ideal for grain yield, proving that fonio could be cultivated in Sudanian and Sudano-Guinean areas. Factor analysis revealed precipitation, C:N ratio, soil pH and texture as the main environmental variables influencing the grain yield in fonio. Our findings contributed to selecting stable, adapted genotypes.

## Introduction

Millets play a critical role in food and nutrition. They are referred to as “hunger crops” with perceived or evidenced attributes such as high resistance to abiotic stresses and superior nutritional value compared to major cereals such as rice, maize and wheat. These crops offer promising solutions to the challenges posed by global population growth, estimated to reach 10 billion by 2050 ^1,2^. Promoting millets in food security programs is crucial for enhancing both agricultural productivity and food security, particularly in developing countries.

Millet crops in West Africa include, for instance, fonio (*Digitaria exilis* (Kippist) Stapf), endemic to the region ^3–5^. It represents an important source of income for smallholder farmers ^6,7^. Cultivated for over 7,000 years, fonio requires low inputs and thrives in degraded and varied soils, and it is particularly adapted to arid and semi-arid regions in sub-Saharan Africa ^8^. Moreover, fonio grains are rich in sulphur-containing amino acids, particularly methionine and cysteine; it is gluten-free, and have a low glycemic index, making them beneficial for people with coeliac disease, obesity and diabetes ^9^. Fonio thus illustrates its potential to contribute to food system diversification and resilience. However, despite these attributes, fonio remains largely an underutilized crop.

Indeed, its production is limited to about eleven West African countries, including Benin ^10^. It is often associated with periods of food scarcity, earning it the nickname “hungry rice” ^11,12^. In Benin, only 813 ha were devoted to fonio production in 2024, yielding 585.69 tons, of which 74% came from Boukoumbe, in the Sudanian phytogeographical area ^10,13^. Benin is one of the countries with very low fonio production, equivalent to only 0.1% of Guinea’s production, the world’s leading producer with 483,006.07 tons in 2024. Compared to other millets such as pearl millet, fonio production remains marginal, representing only about 5% of pearl millet production in Africa in 2024 ^10^. This limited contribution is largely due to the scarcity of improved varieties and the restricted cultivation area ^6,14,15^. This low production, coupled with the increasing population demand, highlights the urgent need to enhance fonio production either through expansion of cultivation area or improvement of its yield.

Over the past 70 years, plant breeding has contributed to more than 50% of yield gains globally ^16^. Plant breeding exploits the genetic diversity of crops to develop or select varieties with improved productivity and high nutritional value ^17,18^. Among the commonly employed methods, assessment of genotype-by-environment interactions is widely used to identify the most productive and stable genotypes across diverse environmental conditions. Interaction between genotype and environment has been explored in the past to identify high grain-yielding genotypes in finger millet, pearl millet and foxtail ^19,20^. Genotype-by-environment interaction (GEI) refers to the differential response of genotypes for a particular trait under varying environmental conditions ^21,22^. When GEI is absent, a single variety could consistently outperform others across all environments ^16,23^. However, in most cases, genotype performance is environment-specific. Among the various analytical tools used to study genotype-environment interactions and genotype performances, the Additive Main Effects and Multiplicative Interaction (AMMI) model and the Genotype Main Effect and Genotype × Environment Interaction (GGE) biplot, and multivariate methods, are the most commonly employed. The AMMI model, developed by Gauch Jr ^24^, combines the Analysis of Variance (ANOVA) with Principal Component Analysis (PCA) to dissect main effects and interactions, thereby providing more precise estimates and performing better than others, such as a linear model. The GGE biplot, which focuses on the combined effects of genotype (G) and GEI simultaneously, complements the AMMI model by offering additional insights into genotype performance across environments and the best-performing or adapted environment ^25^. Those tools also helped to identify stable genotypes by combining multiple traits.

Despite the availability of new genomic resources for fonio, this crop remains poorly improved genetically ^1,3,6^. Several studies have highlighted significant genetic diversity within local fonio resources; nevertheless, officially improved varieties remain scarce or non-existent ^3,26,27^. Genetic improvement efforts are further hampered by botanical constraints, in particular the small size of fonio flowers, which limits the feasibility of controlled crosses ^6,28^. Under these conditions, harnessing existing diversity through selection strategies based on GEI represents a relevant approach. Recent studies indicated that evaluating genotypes across multiple sites is essential to identify stable varieties adapted to diverse environmental conditions ^29,30^. Understanding GEI is therefore a key lever for optimizing fonio production and resilience under climatic constraints. There is also an urgent need to identify the key environmental variables influencing plant performance. It has been demonstrated that several abiotic factors affect the GEI ^16^; among these, soil pH, chemical composition of the soil, temperature, solar radiation, and precipitation are particularly important ^31,32^. Based on these considerations, the following research questions arise: (i) How do genotype by environment interactions influence flowering, maturity, and grain yield of fonio across Benin? (ii) Which genotypes combine high mean performance and stability for these traits across environments? (iii) Are there other cultivable areas that consistently show high performance in fonio production and could therefore serve as target regions for expansion? (iv) Which environmental variables most strongly contribute to genotype-by-environment interactions for flowering, maturity, and yield?

Thus the goals of our study were to (1) evaluate the GEI for days to flowering, days to maturity, and grain yield using AMMI and GGE biplot models, (2) identify the best-performing genotypes across environments for targeted traits, and (3) determine target environments with high potential for fonio expansion, (4) assess the environmental factors influencing GEI under different pedoclimatic conditions. These objectives were addressed through a multilocation trial across various pedoclimatic conditions, with the hypotheses that: (a) significant GEIs exist for all the traits evaluated across the assessment environments; (b) at least one genotype demonstrates both high yield and stability across environments; (c) at least one new environment provides more favourable conditions for higher fonio grain yield compared to traditional environments; (d) environmental factors, such as soil properties and climatic parameters, significantly influence genotype performance and contribute to GEI.

## Methodology

### Study area and plant materials

The experiments were conducted in 2023 across six distinct locations in Benin, each characterized by specific pedoclimatic conditions (Fig. 1, Supplementary Tables S1 and S2). The sites include Boukoumbe and Natitingou (both located within traditional fonio production areas) and Gogounou, Ina, Kandi, and Parakou, which are located outside the main fonio growing areas. These environments are characterized by two phytogeographic areas and are then categorized as followed: Sudanian production area (Boukoumbe and Natitingou), Sudanian non-production area (Gogounou and Kandi) and Sudano-Guinean non-production area (Ina and Parakou). Detailed soil and climatic characteristics of each location are presented in Supplementary Tables S1 and S2. These variations in environmental conditions were intentionally selected to assess genotype performance across a broad range of agroecological contexts. In each location, the trials were hosted either by the National Institute of Agricultural Research of Benin (INRAB) or the University of Parakou to facilitate the implementation and monitoring of field trials.

**Fig. 1.**
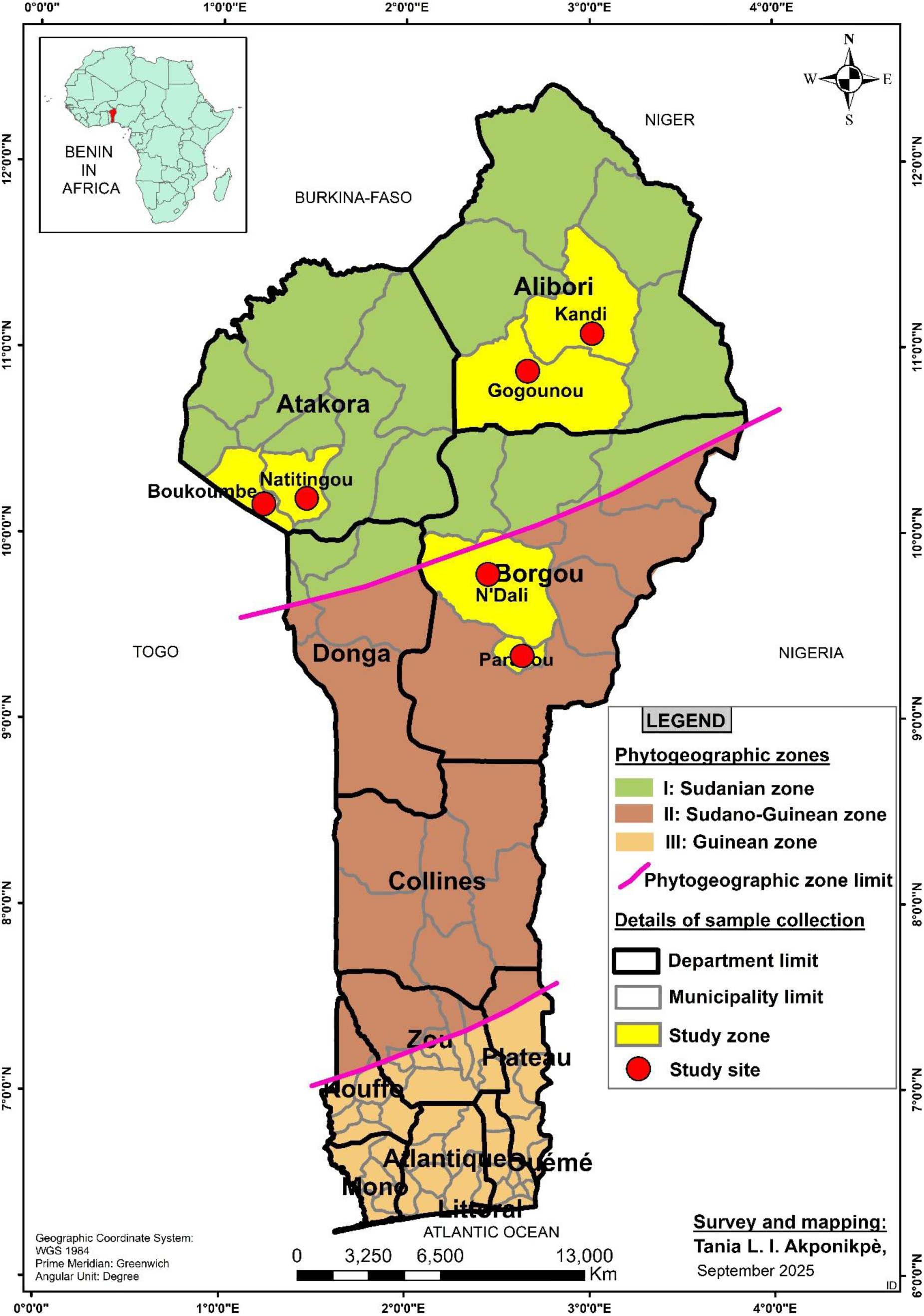
Geographical location of the experimental sites in northern Benin: Boukoumbe, Gogounou, Ina (N’Dali), Kandi, Parakou and Natitingou.

The plant materials consisted of eleven fonio genotypes selected from Ibrahim Bio Yerima et al. ^33^ previous study. This selection was based on agro-morphological and genetic differences (Supplementary Table S3). These genotypes originate from five West African countries: Benin, Niger, Burkina Faso, Mali, and Guinea. Seed accessions were obtained from the Genetics, Biotechnology and Seed Science unit (GBioS) of the University of Abomey-Calavi, where they are conserved in the institutional genebank. These accessions were originally collected by Ibrahim Bio Yerima et al. ^33^.

### Experimental design, management and data collection

The experiments were established at each site using a Randomized Complete Block Design (RCBD) with three replications of eleven genotypes across six environments. Each genotype was sown in 4 m² plots per replicate. Within each plot, rows were spaced 20 cm apart, with planting holes at 20 cm intervals along the rows. Five fonio seeds were sown per hole. Plots were separated by 1 m, and a distance of 1.5 m was maintained between replications. Border rows were established around all trial blocks to minimize edge effects.

Sowing occurred at the onset of the rainy season (June-July 2023) at each site. Weeds were controlled manually by rouging three weeks after planting, followed by regular hand weeding to maintain weed-free plots until harvest. No fertilizers, pesticides, or irrigation were applied during the entire experimentation period to evaluate genotype performance under rainfed, low-input conditions.

Six plants per plot were randomly selected, excluding border plants, to record days to flowering (DTF) and days to physiological maturity (DTM). Grain yield (GRY) was measured based on the total grain weight harvested from each plot and converted to kg ha^-1^.

### Environmental covariates (climate and soil parameters)

Climatic parameters, including mean temperature, relative humidity, total precipitation, and solar radiation during the trials, were obtained from the “Agence Nationale de la Météorologie (METEO-Benin)” of Benin. Prior to the establishment of the experiments, analysis of the collected from each site was carried out to determine key physicochemical properties. Soil texture was assessed using granulometric analysis following the method of Robinson ^34^. Concentrations of major nutrients were determined as follows: total nitrogen by the Kjeldahl method ^35^, available phosphorus by the Bray and Kurtz method ^36^, and exchangeable potassium using the method described by Helmke and Sparks ^37^. Soil pH was measured to evaluate soil acidity, and organic matter content was quantified following the procedure of Bell ^38^. For these analyses, three composite soil samples per site were collected from the 0–20 cm soil layer. Each composite sample was prepared by combining soil collected from the four corners and the center of each replication plot. All soil analyses were performed at the Soil Science Laboratory of the University of Abomey-Calavi.

### Statistical analyses

#### Descriptive statistics and variance components estimation

All statistical analyses were performed using R 4.5.0 software (https://posit.co/download/rstudio-desktop/) ^39^. Descriptive statistics, including means and coefficients of variation, were calculated for each plant variable and genotype within each environment as well as across all environments. We also evaluated trait variation both across and within environments, employing a combined analysis of variance spanning multiple environments:

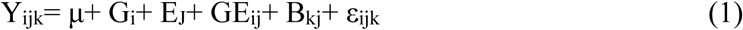

Where Y_ijk_ = observed value of genotype i in block k of environment (location) j, μ = global mean, Gi = effect of genotype i, E_j_ =environment effect, GE_ij_ = the interaction effect of genotype i with environment j, B_kj_ = the effect of block k in environment j, ε_ijk_ = error (residual) effect of genotype i in block k of environment j.

We determined broad-sense heritability H^2^ for each trait studied ^40^, and the genotypic correlation was estimated between the traits for all environments using the software META-R v6.0.4.

#### AMMI model and GGE biplot analysis

The AMMI model, combining the standard analysis of variance with principal component analysis, was used to determine the effect of genotype (G), environment (E) and G × E effects on the traits.

The GGE biplot was used to detect the genotype by environment interaction pattern, classification in mega-environment (which-won-where pattern), and selection of environments based on their ranking compared to an ideal environment.

#### Stability indexes estimation

First, we calculated the AMMI-stability value (ASV) using equation 2 ^41^:

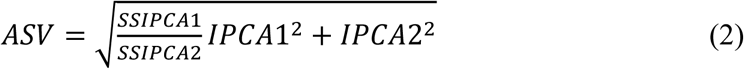

With SSIPCA1 and SSIPCA2, which are the sum of squares of AMMI PC1 and PC2, respectively; IPCA1, the score of the AMMI first component and IPCA2, the score of the AMMI second component. The ASV is the distance from zero in a two-dimensional scattergram of Interaction Principal Component axis 1 (IPCA 1) scores against IPCA 2 scores. Since the IPCA 1 score contributes more to G x E sum of squares, it has to be weighted by the proportional difference between IPCA 1 and IPCA 2 scores to compensate for the relative contribution of IPCA 1 and IPCA 2 to total G x E sum of squares. The lower the ASV, the more stable the genotype.

The second index estimated was the multi-trait stability index (MTSI), according to equation 3

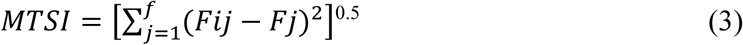

where the MTSI is the multi-trait stability index for the ith genotype, *Fij* is the *jth* score of the *ith* genotype, and *Fj* is the *jth* score of the ideotype. The genotype with the lowest MTSI is then closer to the ideotype.

#### Environmental covariates effect through factorial analysis and correlation

A factor analysis was applied to group environments into factors, and then Pearson correlation was used to assess the effect of environmental covariates (soil properties and the climate parameters) on each of the traits studied. This was computed in R 4.5.0 software ^39^ using the package ASReml and corrplot.

## Results

### Genotypic performance, environmental effect and variance components analysis

Considering both phytogeographic and production areas of fonio, the traits studied varied significantly (p < 0.001) (Fig. 2), except DTF, for which no significant differences were observed. The mean of DTF was consistent across sites, averaging 84.43 days (Fig. 2a). For DTM (Fig. 2b), genotypes matured earlier in the Sudanian non-production area (≈110 days), while longer durations were recorded in the Sudanian production area (117.81 days) and the Sudano–Guinean non-production area (120.13 days). Regarding grain yield (Fig. 2c), the highest mean was recorded at Sudanian non-production area (557.71 kg ha^-1^), followed by the Sudano–Guinean non-production area (380.5 kg ha^-1^), while the lowest yield (101.63 kg ha^-1^) was recorded at Sudanian production area. Notably, both non-production areas significantly outperformed the production area.

**Fig. 2.**
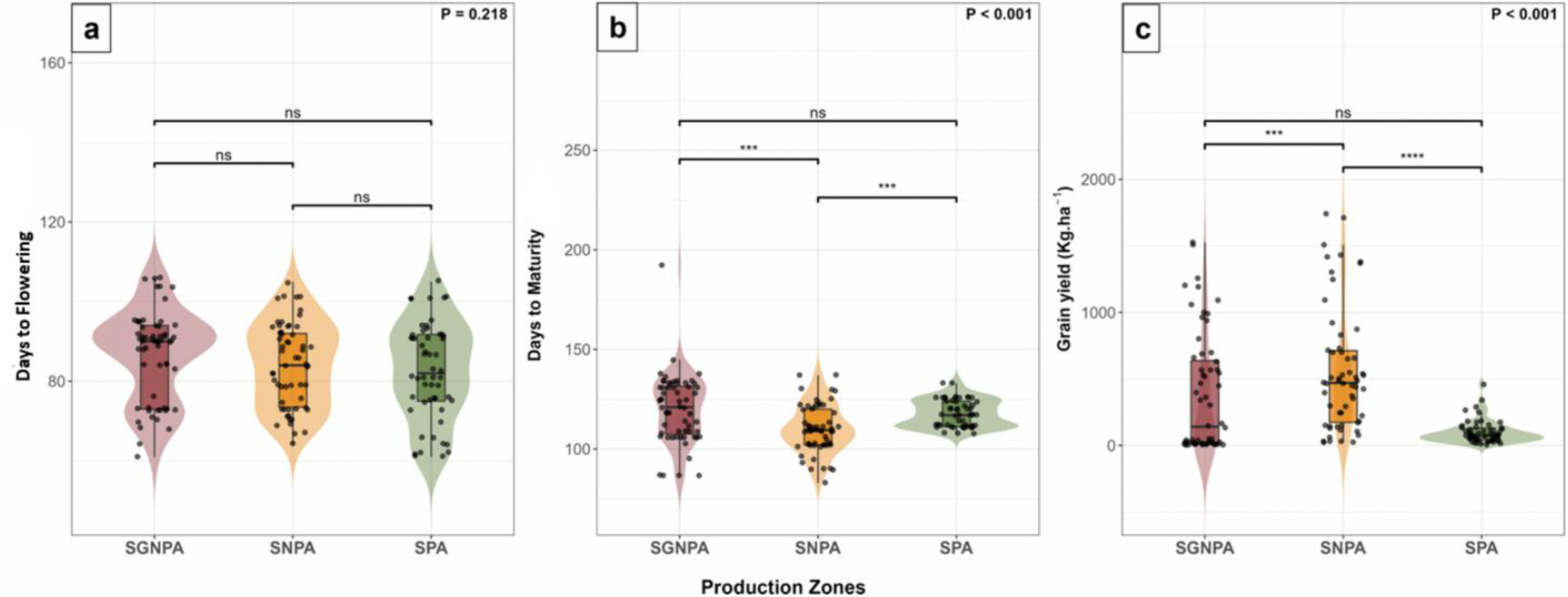
Performance of fonio genotypes in phytogeographical and production areas. **a:** days to flowering (**DTF**), **b:** days to maturity (**DTM**), and **c:** grain yield (**GRY** in kg ha^-1^). **SGNPA:** Sudano-Guinean non-production area, **SNPA:** Sudanian non-production area, **SPA:** Sudanian production area. ***: the significance levels at p < 0.001, **ns**: non-significant at p > 0.05.

No significant differences were observed among phytogeographic and production areas for genotypes B12, B33, B37, BF1, G31, and G32 across the assessed traits (Figs. 3-1 to 3-3). Their mean grain yields across the three phytogeographic and production areas were 485.35, 314.60, 238.85, 433.94, 463.02 and 438.56 kg ha^-1^, respectively (Fig. 3-3). Interestingly, the highest yields for these genotypes were recorded in non-production areas. Notably, genotypes B12 and G31 achieved average yields of 817.4 and 737.27 kg ha^-1^, respectively, in the Sudanian non-production area, while B33 reached 634.44 kg ha^-1^ in the Sudano–Guinean non-production area (Fig. 3-3). Only genotype M14 showed significant environmental variation for GRY, achieving its highest performance of 597.35 kg ha^-1^ in the Sudanian non-production area. This genotype also demonstrated significant differences (p < 0.05) in DTM across environments (Fig. 3-2), requiring an average of 97.2 days in the Sudanian non-production area compared to 100.67 days in the Sudano-Guinean non-production area and 111.17 days in the Sudanian production area. Regarding this trait, genotypes G4, M15, M5, and N35 also showed significant differences across the three phytogeographic and production areas, with the Sudano-Guinean non-production area consistently required fewer days to maturity. For DTF (Fig. 3-1), significant environmental differences were observed only for genotypes G4 and N35, both requiring approximately 79 days in the Sudanian non-production area, which was lower than in the other environments.

**Fig. 3-1.**
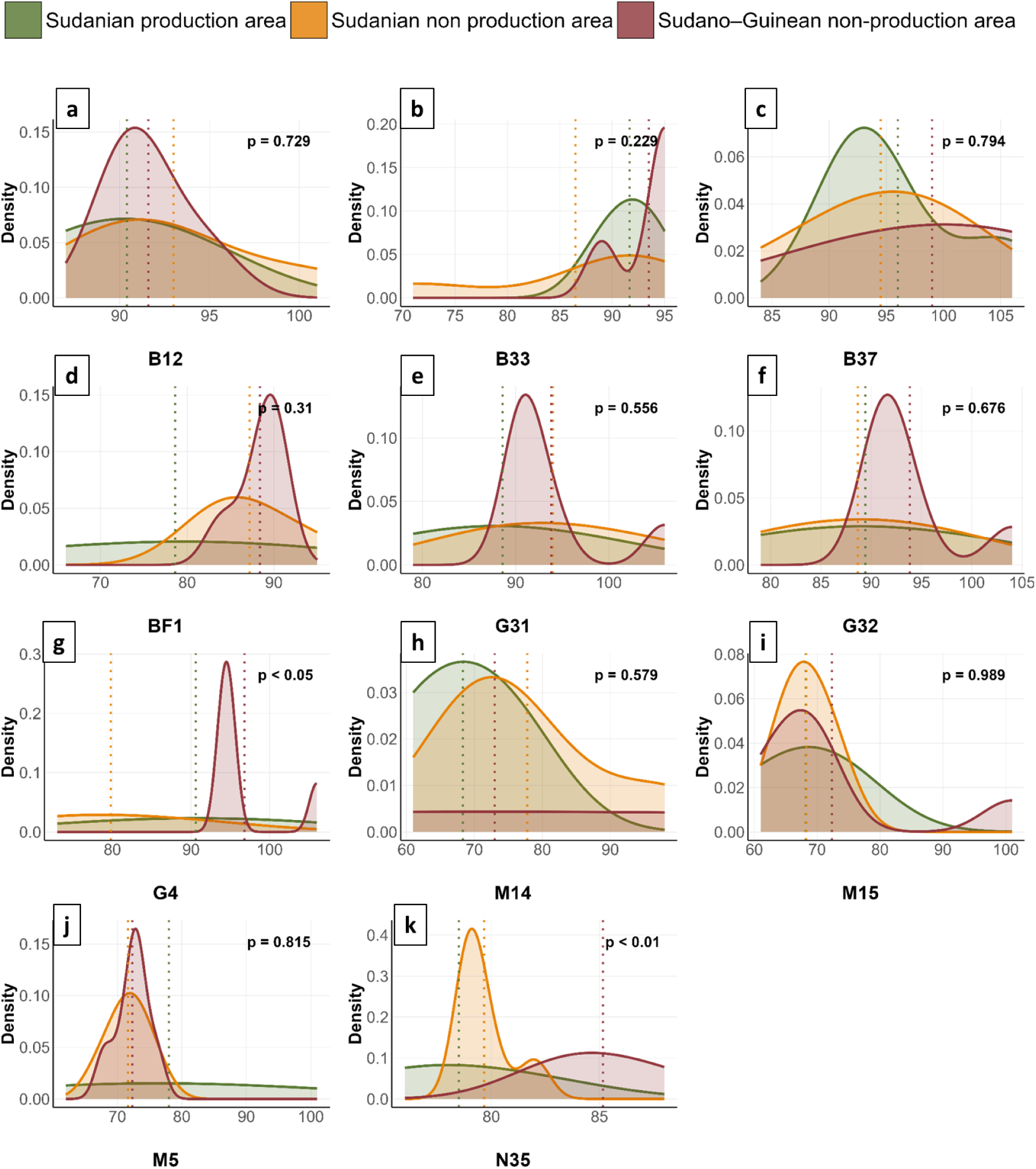
Distribution of phenotypic values for days to flowering (DTF) among phytogeographical and production areas of eleven genotypes.

A high level of heritability in the broad sense was observed across all traits and environments (0.98 for DTF, 0.94 for DTM, and 0.74 for GRY). Genetic correlations between the traits studied were 0.57 between DTM and GRY, 0.66 between DTF and GRY, and 0.99 between DTF and DTM (Table 1). These positive correlation values thus indicate genetic similarity between DTM and GRY on the one side, and DTF and GRY on the other. Very high genetic similarity was recorded between DTF and DTM.

**Table 1.** Genetic parameters for days to flowering (DTF), days to maturity (DTM) and grain yield (GRY).

| Variables | Broad-sense heritability (H<sup>2</sup>) | Selection accuracy (SA) | Genetic correlation |  |
| --- | --- | --- | --- | --- |
|  |  |  | DTM | DTF |
| Days to flowering (DTF) | 0.98 | 0.99 | - | - |
| Days to maturity (DTM) | 0.94 | 0.97 | - | 0.99 |
| Grain yield (GRY) | 0.74 | 0.86 | 0.57 | 0.66 |

### AMMI model for analysing genotype performance and stability

Table 2 shows the AMMI model for DTF, DTM, and GRY. For the DTF trait, a highly significant effect (p < 0.001) of genotype, environment and GEI, which explained respectively 47.4%, 9.4% and 14.9% of the total variation, was observed. The DTM trait was significantly affected by environment and genotype (p < 0.001), with respective variances of 17.2% and 32.5%. Variation due to the GEI (14.5%) was also significant at p < 0.05 for this same trait. For the grain yield, a significant effect (p < 0.001) of environment and genotype was observed, explaining 53.5% and 6.5% of the total variation, respectively; 15.6% of the yield variation was due to GEI with a significant effect at p < 0.01.

**Table 2.** AMMI model for days to flowering (DTF), days to maturity (DTM) and grain yield (GRY).

|  | Df | Days to flowering (DTF) |  |  | Days to maturity (DTM) |  |  | Grain yield (GRY) |  |  |
| --- | --- | --- | --- | --- | --- | --- | --- | --- | --- | --- |
|  |  | SS | MS | %SS | SS | MS | %SS | SS | MS | %SS |
| **Environment (ENV)** | 5 | 2583 | 516.5*** | 9.41 | 6390 | 1278*** | 17.22 | 16517679 | 3303536*** | 53.48 |
| **Rep (ENV): Design** | 12 | 432 | 36 | 1.57 | 612 | 51.00 | 1.65 | 2010937 | 167578*** | 6.51 |
| **Genotype** | 10 | 13020 | 1302*** | 47.45 | 12077 | 1207*** | 32.54 | 2019641 | 201964*** | 6.54 |
| **G x E** | 50 | 4106 | 82.1*** | 14.96 | 5398 | 108* | 14.54 | 4827082 | 96542** | 15.63 |
| **Residual** | 94 | 2216 | 23.6 | 8.08 | 6832 | 72.7 | 18.41 | 49415 | 52569.00 | 0.16 |
| **PC1** | 14 | 2003 | 143.1*** | 7.3 | 3473.6 | 248.1*** | 9.36 | 2938964 | 209926*** | 9.51 |
| **PC2** | 12 | 1311 | 109.2*** | 4.78 | 1175.8 | 98.0 | 3.17 | 1344928 | 112077*** | 4.35 |
| **PC3** | 10 | 905 | 90.5*** | 3.3 | 613.6 | 61.4 | 1.65 | 1122545 | 112255* | 3.63 |
| **PC4** | 8 | 516 | 64.5** | 1.88 | 458.6 | 57.3 | 1.24 | 32426 | 4053* | 0.1 |
| **PC5** | 6 | 349 | 58.2* | 1.27 | 83.8 | 14.0 | 0.23 | 24658 | 4110.00 | 0.08 |

**Df:** degrees of freedom, **SS:** sums of squares, **MS:** mean square, **%SS:** proportion of SS associated with each factor over the total SS, **ENV:** environment, **Rep:** replication, **G x E:** Genotype by Environment interaction, **PC:** principal component. \*, \*\* and \*\*\* indicate the significance levels at $p < 0.05$ , $0.01$ and $0.001$ , respectively, and ns indicates non-significant at $p > 0.05$ .

Figures 4a to 4c show the biplot of the additive main effects and multiplicative interaction (AMMI) between the PC1 axis and the DTF, DTM, and GRY traits. The PC1 axes explains 39.4%, 59.8% and 53.8% of the observed variance respectively for DTF, DTM and GRY. The x-axes shows the traits studied values. The average DTF was around 85 days and 116 days for DTM. M15, M5, and M14 genotypes exhibited early flowering and maturity, whereas B37, B12, and G31 genotypes were late. Early flowering was observed at the Natitingou and Gogounou sites, while early maturity at only Gogounou site (Figs. 4a and 4b). An average grain yield of around 400 kg ha^-1^ was recorded (Fig. 4c). Ina and Gogounou had the highest grain yield and the best-performing genotypes were G31, N35, BF1, G32, and B12.

**Fig. 4-2.**
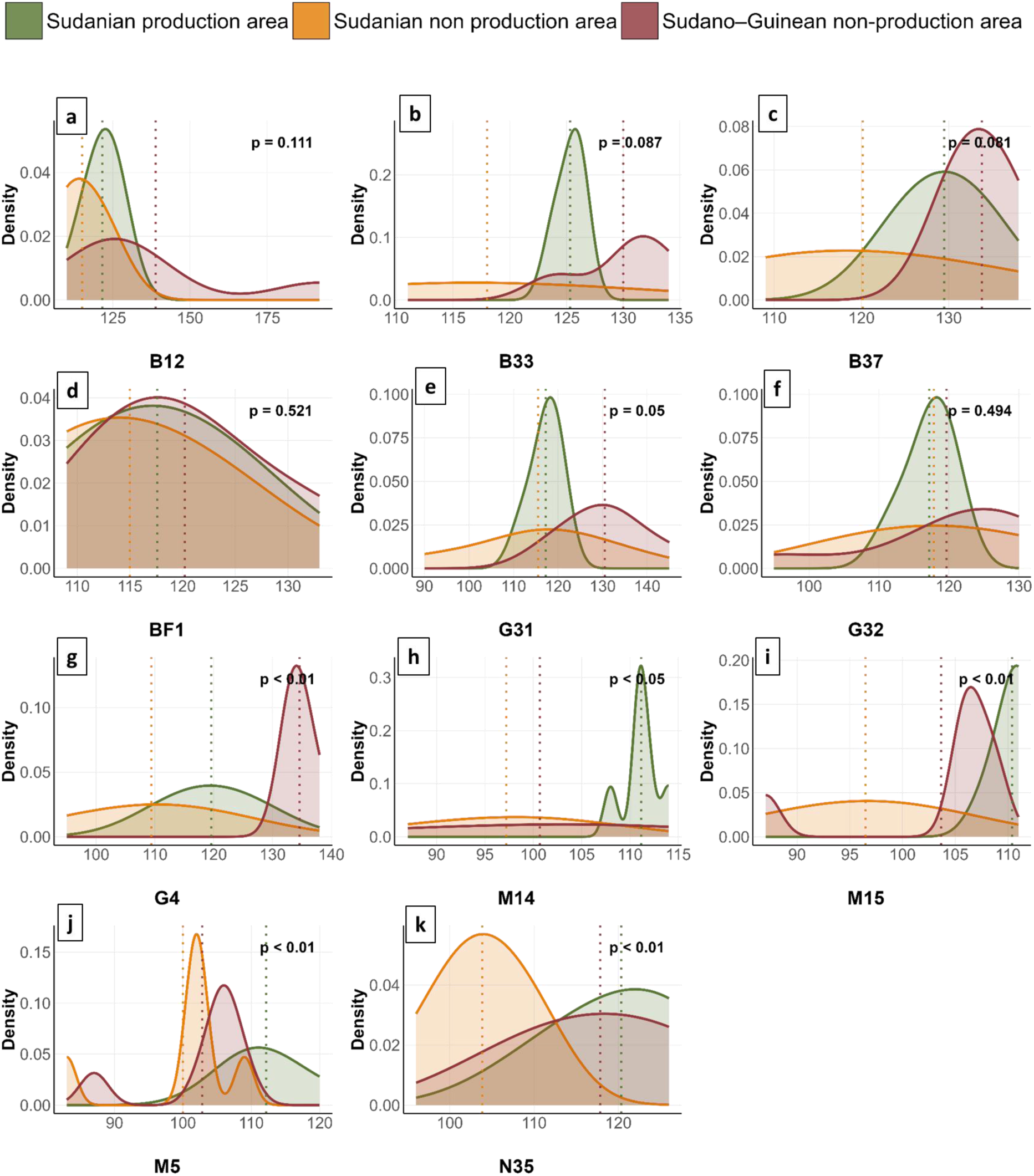
Distribution of phenotypic values for days to maturity (DTM) among phytogeographical and production areas of eleven genotypes.

Figures 4d to 4f show the interaction of the first two principal components of the AMMI 2 biplot model, which accounted for 65.2%, 80.1%, and 78.4% of the total variation in the sums of squares of the GEI for DTF (Fig. 4d), DTM (Fig. 4e) and GRY (Fig. 4f), respectively. Genotypes (black colour) and environments (red colour) located far from the biplot origin showed the highest effects of GEI, whereas genotypes and environments close to the origin showed the lowest influence of GEI. Thus, Ina and Boukoumbe, with their short vector, did not exert strong interactive forces for DTF and GRY; Parakou also had a short vector for the DTM trait, so it exerted a slow interactive force. Similarly, genotypes N35, M15, and M5 were stable and less sensitive to the interaction for DTF; B33 and B37 were less sensitive to interaction for DTM, and G31 and B37 were the most stable genotypes for grain yield, explained by their proximity to the origin.

### GGE biplots analysis

#### Which-won-where pattern for selecting adapted genotypes across environments

Figures 4g to 4i show GGE biplots displaying the interaction among the eleven fonio genotypes and the six environments for DTF, DTM, and GRY. The polygon view of the GGE biplot pattern revealed 88.7% for DTF (Fig. 4g), 90.4% for DTM (Fig. 4h), and 81.9% for GRY (Fig. 4i) of the variation. There were 6, 5, and 6 biplot sections for DTF, DTM, and GRY, respectively. Within these sections, 2, 2, and 3 mega-environments were observed for DTF, DTM and GRY, respectively, each with a winning genotype. A mega-environment represents a group of environments clustered within the same biplot section.

For DTF (Fig. 4g), the first mega-environment comprised Gogounou and Natitingou, and the second Boukoumbe, Kandi, Parakou and Ina. The vertex polygon included genotypes B37, B33, M14, M15, M5 and G4, representing high-value genotypes. For this trait, genotypes with high values were late-flowering. The vertex genotype B33 was late-flowering in the first mega-environment, while B37 and G4 were late-flowering genotypes in the second mega-environment. Genotypes M5, M14, and M15, which were not found in the mega-environments, recorded the lowest values across the environments studied and were therefore early-flowering. In Fig. 4h, representing DTM, the vertex polygon was formed by B37, B12, M15, M14, and G32 and had high values, indicating late maturity. The two mega-environments were distributed as follows: (1) Ina and (2) Boukoumbe, Gogounou, Kandi, Natitingou, and Parakou. The points representing the two mega-environments were very close, reflecting their similarity in this trait. For the first mega-environment, B12 was a late-maturing genotype, while B37 was late in the second mega-environment. The polygon’s other genotypes (M14 and M15), which were not found in the mega-environment sections, were therefore early-maturing genotypes irrespective of environment. For the GRY (Fig. 4i), the first mega-environment included Natitingou, Boukoumbe, and Kandi, the second mega-environment included Ina, Gogounou and the third only Parakou. The vertices of the polygon were composed of the N35, G32, B37, M15 and M5 genotypes. Genotype N35, being the vertex of the first mega-environment, was therefore the most productive in this mega-environment. Genotype G32 had the highest value in the second mega-environment, while M15 showed the highest value in the third mega-environment. Genotypes B37 and M5 not appearing in any mega-environment were considered low-performance genotypes regardless of environment.

#### Ranking environment from GGE biplot: ideal environment assessment

Figures 4j to 4l represent the ranking of environments according to each trait studied. For DTF and DTM, a closed environment to the biplot origin correspond to delayed flowering and maturity. For these traits, the ideal environment is located farthest from the origin; therefore, Gogounou and Boukoumbe were the optimal environments for DTF (Fig. 4j) and DTM (Fig. 4k). Ina was the most desirable for GRY (Fig. 4l). In contrast, Parakou was the least desirable environment for GRY.

### Stable genotype identification through stability indexes

ASV, AMMI-stability value indexes are presented in Supplementary Table S5 for each trait and their rank (from most stable to least stable). The M15, N35, and M5 genotypes were the most stable, for the DTF trait. In all environments, these stable genotypes showed earliness at flowering (M15 = 69.87 days; N35 = 81.1 days; M5 = 73.77 days). For DTM, the most stable genotypes were B37, B33 and BF1, with respective averages of 127.2, 124.36 and 117.60 days, among the highest averages and therefore the latest to reach maturity. The fourth most stable genotype, M15, with 104 days to maturity, was one of the earliest genotypes. G31, M14 and B37 were the most stable in grain yield, with 480.28, 262.35 and 268.26 kg ha^-1^. G31 was one of the highest-performing genotypes in all environments; N35 and B12 were also medium-high-performing genotypes, but were less stable.

The Multi-Trait Stability Index (MTSI) was used for simultaneous selection for DTF, DTM, and GRY among the genotypes. The selection differential was negative for yield. The selection differential for the Weighted Average of Absolute Scores (WAASB) index was 56.4%, 49.6%, and 34.6% for DTF, DTM, and GRY, respectively (Supplementary Table S6). Figure 5 shows two selected genotypes with the lowest MTSI values: M5 and M15, which represent the most stable genotypes.

**Fig. 5-3.**
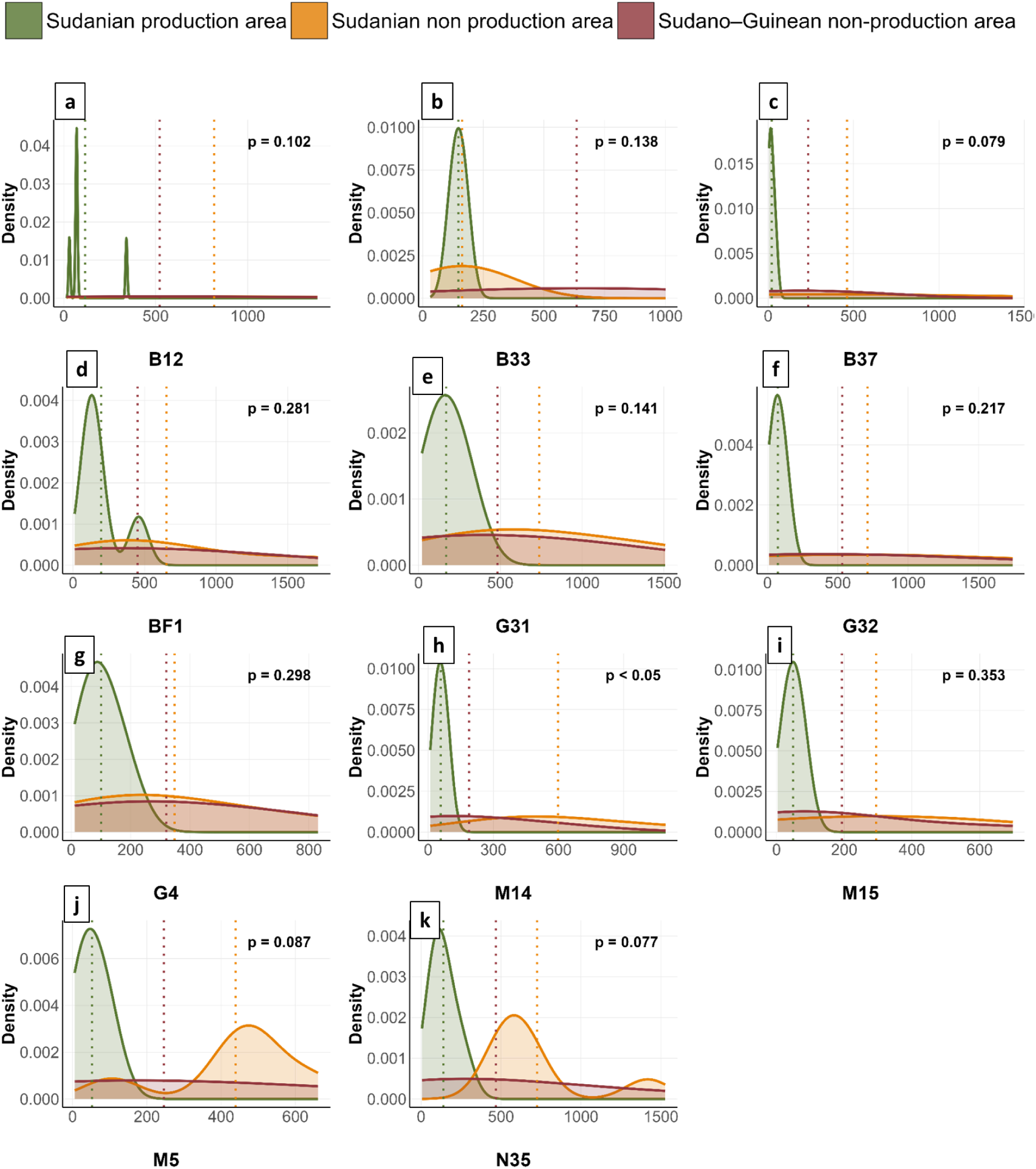
Distribution of phenotypic values for grain yield (kg ha^-1^) among phytogeographical and production areas of eleven genotypes.

**Fig. 6.**
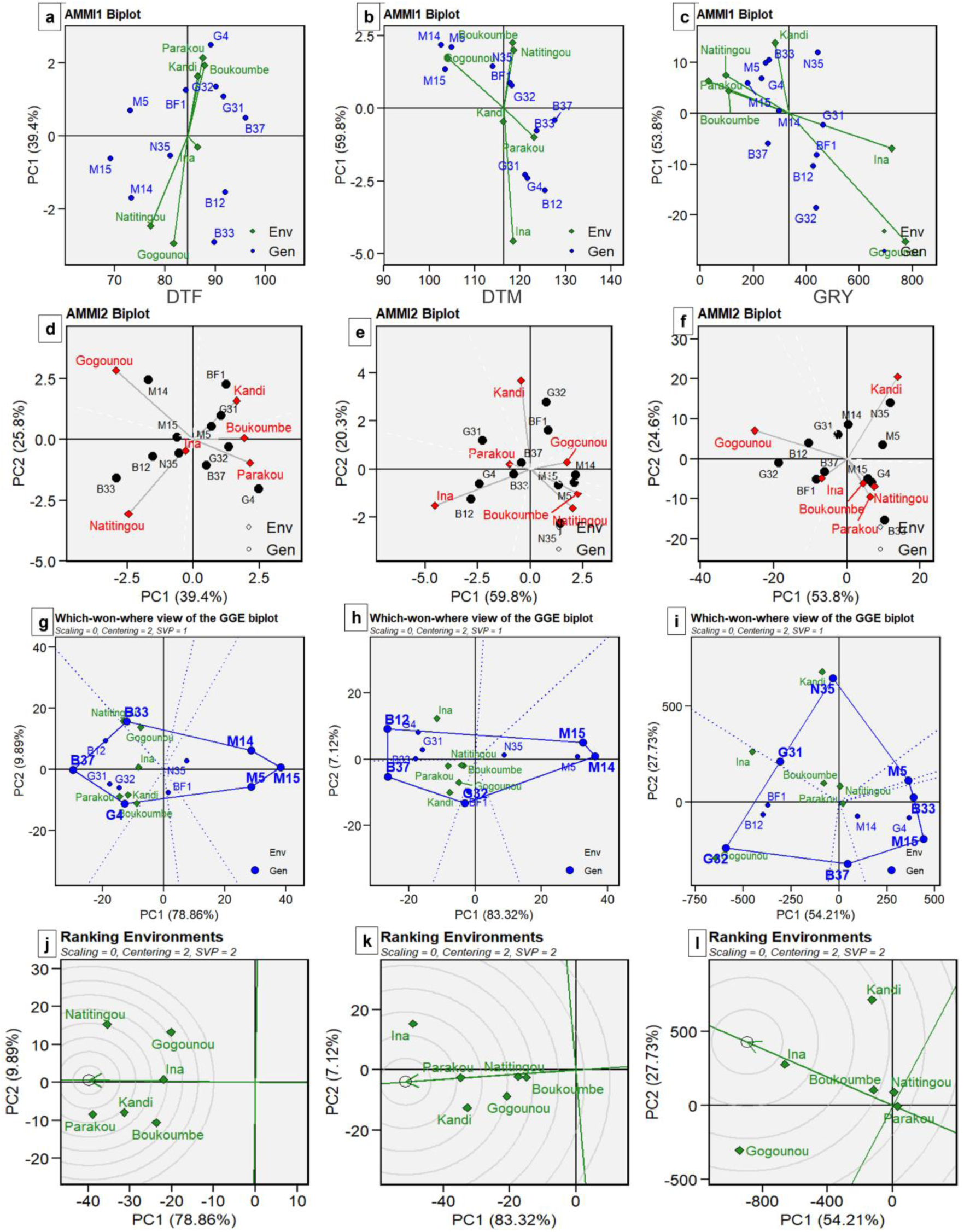
Additive main effects and multiplicative interaction (AMMI) and genotype main effect plus genotype by environment interaction effect (GGE) biplots model. AMMI 1 biplot for a: DTF, b: DTM, c: GRY. AMMI2 biplot of the first two principal components (PC1 and PC2) for d: DTF, e: DTM, f: GRY. (This involves projecting the variance of the interaction of the eleven genotypes in the six environments onto the first two axes. The centre of the biplot (0, 0) was divided into four distinct sections passing through the two lines, vertically and horizontally)**. “Which-won-where” pattern of GGE biplot for g: DTF, h: DTM, i: GRY. Ranking environments of GGE biplot; j: DTF, k: DTM, l: GRY (**The centre of the set of circular lines represents the “ideal” environment, allowing us to measure the distance between an environment and the ideal environment)**. Env:** Environment**, Gen:** Genotype**, DTF:** days to flowering**, DTM:** days to maturity and **GRY:** grain yield (kg ha^-1^).

**Fig. 7.**
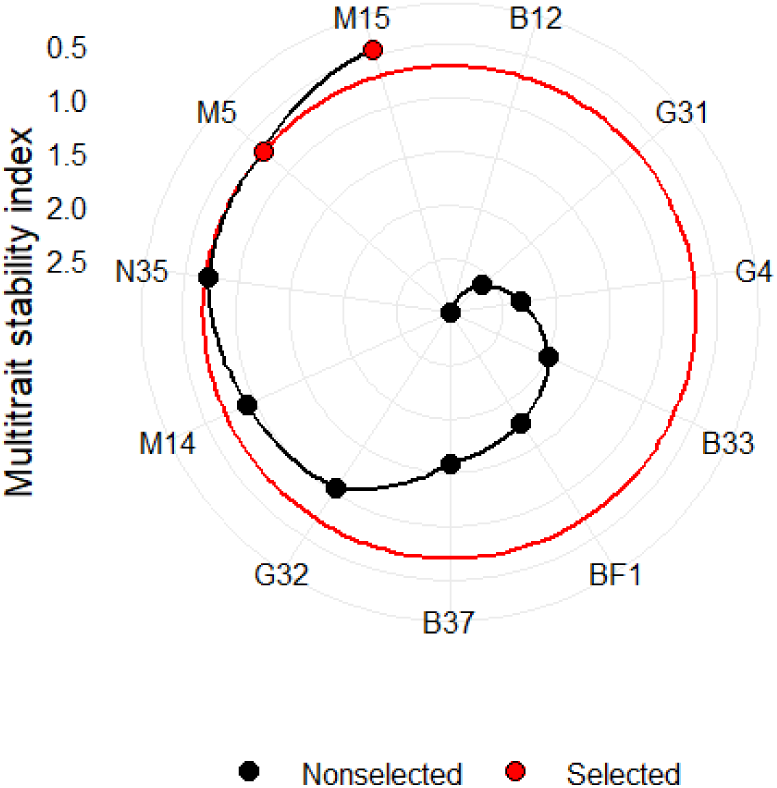
Fonio genotypes selected by multi-trait stability index based on days to flowering, days to maturity and grain yield.

### Environmental factors influencing G x E: Factor analysis and correlation between environmental covariates and loading factors

Supplementary Table S7 illustrates the contribution of each environment to its respective factor. For DTF and DTM, all environments studied loaded primarily on FA1, indicating similar environmental influences. However, for GRY, the environments were separated into two groups: Boukoumbe, Gogounou and Ina were grouped in FA1, and Kandi, Natitingou and Parakou were in FA2.

In Table 3, the Pearson correlation coefficients between environmental loadings and environmental covariates are presented for the two factors. From these results, each environmental covariate effect on the responses of the environments was assessed, identifying the factors most likely to influence the GEI. Therefore, the overall precipitation affected genotype performance in environments grouped by FA1 for DTF (Table 3). Considering DTM, environments clustered in the FA1 were most characterized by soil organic matter, C: N ratio and total nitrogen, solar radiation and the sand, silt and clay proportion, representing the most influential factors in these environments. Finally, for GRY, overall precipitation and clay proportion were the most important factors of the environments grouped in FA1. While soil pH, C: N ratio, CEC, and phosphorus content strongly influenced environments grouped in FA2 for the grain yield.

**Table 3.** Pearson’s correlation coefficient between the environmental covariates and the environmental loadings extracted from the factor analysis.

|  | Days to flowering<br>(DTF) |  | Days to maturity<br>(DTM) |  | Grain yield (GRY) |  |
| --- | --- | --- | --- | --- | --- | --- |
|  | FA1 | FA2 | FA1 | FA2 | FA1 | FA2 |
| Average temperature (°C) | 0.03 | -0.31 | -0.02 | -0.67 | -0.01 | -0.14 |
| Overall precipitation (mm) | **-0.56** | 0.02 | -0.22 | 0.69 | **0.39** | 0.00 |
| Solar radiation (MJ/m<sup>2</sup>/j) | -0.10 | -0.30 | -0.41 | -0.47 | -0.07 | -0.26 |
| Relative Humidity (%) | -0.07 | 0.46 | 0.15 | 0.45 | 0.06 | 0.16 |
| Clay (%) | 0.13 | -0.16 | **-0.77** | 0.76 | **-0.42** | -0.09 |
| Silt (%) | -0.32 | 0.12 | **-0.77** | 0.67 | -0.10 | -0.15 |
| Sand (%) | 0.19 | -0.03 | **0.81** | -0.73 | 0.21 | 0.13 |
| Soil pH | 0.06 | -0.65 | -0.19 | 0.55 | 0.23 | **-0.84** |
| Organic carbon (%) | 0.00 | -0.04 | **-0.81** | 0.81 | -0.31 | -0.23 |
| Total Nitrogen (%) | 0.04 | -0.09 | **-0.63** | 0.84 | -0.31 | 0.04 |
| C:N ratio | -0.25 | -0.06 | **-0.80** | 0.55 | -0.03 | **-0.63** |
| Cation exchange capacity (meq/100) | 0.19 | -0.25 | 0.14 | 0.73 | -0.04 | 0.30 |
| Phosphorus (ppm) | -0.22 | -0.26 | -0.14 | -0.60 | 0.24 | **-0.59** |

Note. The highest correlations between environmental covariates and the environmental loadings of the factorial model are highlighted in bold. **FA1:** Factor 1, **FA2:** Factor 2.

## Discussion

The performance of eleven fonio genotypes was evaluated in a multi-environment trial across six different locations in northern Benin, involving both traditional and non-traditional production areas with diverse pedoclimatic conditions. The objectives were to better understand the genotype × environment interaction (GEI) for three agronomic traits, identify environments for fonio expansion and assess pedoclimatic factor impacts on fonio production.

The AMMI analysis highlighted the strong influence of environmental conditions, which played a limited role in shaping phenological traits (DTF and DTM), but exerted a major effect on grain yield. This was also confirmed by the significant differences observed among phytogeographic and production areas. Similar results have been reported for finger millet, little millet, tef, foxtail millet and maize ^20,43–47^, indicating the influence of edaphic and climatic factors, often identified as determining covariates in rainfed systems. In contrast, genotype effects were predominant for phenological traits (47% for DTF and 33% for DTM), suggesting strong control. This was supported by high broad-sense heritability values for DTF (0.98) and DTM (0.93), compared to a lower value for GRY (0.74). High heritability for phenological traits indicates that genetic factors outweigh environmental influences, making them reliable targets for breeding programs ^20^. The pronounced environmental effect on GRY may reflect differences in nutrient use efficiency among genotypes ^14^ or their specific adaptation to local conditions. For instance, early flowering in genotypes from arid regions helps in drought-escape strategy ^4,48^, a pattern also observed in this study with genotypes from Mali. In contrast to precocity to flowering and maturity, grain yield was mainly driven by environmental variation, as reported in tef and foxtail millet, crops close to fonio ^44,49^. In studies conducted in Northern Benin, a similar trend in environment and genotype effects as well as their interaction was also reported in sesame accessions, with the environment accounting for the largest share of the total variation^50^. The significant GEI observed implies specific adaptation of certain genotypes to particular environments ^51^. Grouping of environments into the same mega-environments reflects strong positive correlations in genotype performance across sites ^52,53^. Thus, the most productive genotypes in these groups were N35, G32, and M15, respectively.

Here, traditional fonio production areas (Boukoumbe and Natitingou) and no fonio production areas (Ina, Gogounou, Parakou and Kandi) have been selected in different phytogeographic areas (Sudanian and Sudano-Guinean). The environments were then categorized as Sudanian production area, Sudanian non-production area and Sudano-Guinean non-production area. In general, the Sudanian non-production and Sudano–Guinean non-production areas exhibited superior performance in terms of DTM and GRY. This may be attributed to its pedoclimatic conditions, which likely favor optimal fonio development. Fonio performs best under moderate rainfall, well-drained soils, and relatively stable temperatures^54^. These conditions appear to align with those found in the Sudanian non-production area. The Sudano-Guinean area was reported to be the preferred climate of fonio crop^55^. Nevertheless, fonio in Benin is traditionally recognized as being predominantly cultivated in the Atacora department, within the Sudanian production area characterized by a mountainous environment, low rainfall and poor soils ^55^. Surprisingly, the lowest grain yields were recorded across the tested genotypes in this area. This paradox raises the possibility that environmental conditions in the natural production area are becoming less favourable for fonio. One explanation could be the effects of climate change, with increased heat or extreme cold events in mountainous areas reducing crop productivity ^56,57^. In the same perspective, the GGE biplot pattern, ranking plot, has identified Ina (Sudano-Guinean area) and Kandi (Sudanian) as ideal environments for high-yield fonio cultivation. This confirms opportunities to expand production beyond traditional areas. Ina’s high performance for GRY is associated with moderate rainfall, low clay content, and low nutrient levels, consistent with optimal conditions ^54^. Factor analysis confirmed that precipitation, soil texture, soil pH and C: N ratio were the main environmental variables influencing the grain yield in fonio. This result is consistent with earlier studies ^16,58–60^. Moreover, great contribution of precipitation in grain yield variation for Kersting’s groundnut in Benin was earlier reported ^61^. It would be very interesting to know better the ideal ecological parameters for fonio crop production, like for major cereals.

According to the AMMI stability value and AMMI1, genotypes M15, N35, and M5 exhibited the highest stability for days to flowering, with M15 and M5 also being early-flowering. For days to maturity, genotypes B37, B33, and BF1 were the most stable, but B37 and B33 were late-maturing genotypes. However, this stability was associated with late maturity rather than early maturity. Regarding grain yield, genotypes G31, M14, and B37 showed the greatest stability, with G31 combining stability and high yield. The GGE biplot analysis corroborated these stability patterns. Furthermore, the Multi-Trait Stability Index (MTSI), developed by Olivoto et al. ^42^, identified M5 and M15 as stable across all traits, but did not have the highest yield, illustrating that stability is valuable only when combined with strong performance ^62^. Overall, the instability of yield across environments reflects the strong influence of environmental variability, as also reported by Kang ^16^. Validation in additional locations and across multiple seasons will be crucial to strengthen these findings. Furthermore, involving farmers in the genotype selection process will help ensure that the selected genotypes align with their preferences, thereby increasing the likelihood of adoption in rural areas. Combining plant breeding tools with farmers’ participation is a consistent and effective approach for further utilization ^63,64^. Moreover, our findings indicate that the suitability of traditional production areas for fonio in Benin may be shifting. Despite its historical and cultural importance, the Sudanian production area may no longer provide optimal conditions for stable fonio yields. This highlights the urgent need to reassess production strategies and to integrate climate and soil considerations into future breeding and cultivation programs. Nevertheless, by testing a wide range of environments, this study has identified promising new production areas for fonio, potentially expanding its cultivation beyond the Atacora region.

## Conclusion

This multi-environment evaluation of eleven fonio genotypes in northern Benin revealed substantial phenotypic variability, with phenological traits such as days to flowering and days to maturity under strong genetic control and grain yield (GRY) mainly influenced by environmental factors. The significant genotype × environment interactions observed highlight both the potential for identifying broadly adapted and stable genotypes and the need for site-specific recommendations. The identification of three distinct mega-environments and the unexpected high performance of fonio at certain non-traditional production areas, such as Ina, demonstrate opportunities for expanding fonio cultivation beyond its current range. These findings provide a scientific basis for targeted breeding, the selection of adapted genotypes, and the promotion of fonio as a resilient, underutilized crop contributing to food security in West Africa. Future work should integrate multi-year trials, advanced genomic tools, more genotypes and farmer participatory approaches to enhance adoption and impact.

## Data availability

The data that support the findings of this study are available from the first author upon reasonable request.

## Figure caption

**Fig. 8. Geographical location of the experimental sites in northern Benin: Boukoumbe, Gogounou, Ina (N’Dali), Kandi, Parakou and Natitingou.**

**Fig. 9. Performance of fonio genotypes in phytogeographical and production areas. a:** days to flowering (DTF), b: days to maturity (DTM), and c: grain yield (GRY in kg ha-1). SGNPA: Sudano-Guinean non-production area, SNPA: Sudanian non-production area, SPA: Sudanian production area.

**Fig. 10-1. Distribution of phenotypic values for days to flowering (DTS) among phytogeographical and production areas of the eleven studied genotypes.**

**Fig. 11-2. Distribution of phenotypic values for days to maturity (DTM) among phytogeographical and production areas of the eleven studied genotypes.**

**Fig. 12-3. Distribution of phenotypic values for grain yield (kg ha**^-1^**) among phytogeographical and production areas of the eleven studied genotypes.**

**Fig. 13. Additive main effects and multiplicative interaction (AMMI) and genotype main effect plus genotype by environment interaction effect (GGE) biplots model. AMMI 1 biplot for a: DTF, b: DTM, c: GRY. AMMI2 biplot of the first two principal components (PC1 and PC2) for d: DTF, e: DTM, f: GRY. “Which-won-where” pattern of GGE biplot for g: DTF, h: DTM, i: GRY. Ranking environments of GGE biplot; j: DTF, k: DTM, l: GRY. Env:** Environment, **Gen:** Genotype, **DTF:** days to flowering, **DTM:** days to maturity and **GRY:** grain yield (kg ha^-1^).

**Fig. 14. Fonio genotypes selected by multi-trait stability index based on days to flowering, days to maturity and grain yield.**

## Acknowledgements

Tania L. I. Akponikpè’s PhD is a scholar of the University of Abomey-Calavi (Republic of Benin) and the Université de Lorraine (France). Her research work was funded by French National Research Agency through the PEA-BIOVALOR (ANR-21-PEA2-0006) Biovalor project. We sincerely thank the “Institut National des Recherches Agricoles du Bénin” (INRAB) and La Maison du Fonio for their valuable support in establishing the experiments. We are also grateful to our dedicated field assistants (Hadid A. Gangni-Ahossou, Hunic Akodjenou, Monas Lonou, Caleb Sacca, Farid Guissida, and Joël O. Sossoukpe Alido) for their commitment during trial implementation and data collection.

## Author contributions

**T.L.I.A.:** Data curation, Formal analysis, Investigation, Methodology, Software, Visualization, Writing – original draft, Writing – review and editing**, E.L.S.:** Conceptualization, Data curation, Supervision, Writing – review and editing**, I.A:** Formal analysis, Software, Visualization, Writing – original draft, Writing – review and editing**, A.R.I.B.Y.:** Investigation, Methodology, Writing – review and editing, **S.P.:** Conceptualization, Data curation, Supervision, Writing – review and editing**, G.L.A.:** Conceptualization, Data curation, Supervision**, E.G.A-D.**: Conceptualization, Data curation, Funding acquisition, Methodology, Project administration, Resources, Supervision, Validation, Writing – review and editing.

