## Supplementary files for "Genotype-by-environment interaction analysis for flowering, maturity time and yield in fonio across traditional and prospective production areas in Northern Benin"

**Supplementary information**

**Table S1. Climatic characteristics of the study area**

| **Environments** | **Latitude** | **Longitude** | **Mean temperature (°C)** | **Overall precipitation (mm)** | **Solar radiation (MJ/m^2^/j)** | **Relative Humidity (%)** |
| --- | --- | --- | --- | --- | --- | --- |
| **Boukoumbe** | 10.22 | 1.11 | 26.03 | 1099.32 | 18.05 | 82.74 |
| **Gogounou** | 10.74 | 2.81 | 26.85 | 993.88 | 18.52 | 76.39 |
| **Ina** | 9.96 | 2.70 | 25.93 | 1111.84 | 17.07 | 83.18 |
| **Kandi** | 11.33 | 2.99 | 27.95 | 689.74 | 19.3 | 70.14 |
| **Natitingou** | 10.15 | 1.40 | 26.45 | 1005.4 | 18.2 | 76.03 |
| **Parakou** | 9.31 | 2.71 | 25.88 | 893.17 | 17.07 | 83.89 |
| **Source:** METEO-Benin. | | | | | | |

**Table S2. Average values of measured soil properties of the study area**

| **Environments** | **Clay (%)** | **Silt (%)** | **Sand (%)** | **pH** | **Organic carbon (%)** | **Total N (%)** | **C/N** | **CEC (meq/100)** | **Assimilable phosphorus (ppm)** |
| --- | --- | --- | --- | --- | --- | --- | --- | --- | --- |
| **Boukoumbe** | 11.97 | 28.80 | 58.64 | 5.19 | 1.14 | 0.10 | 11.79 | 7.5 | 20.66 |
| **Gogounou** | 9.80 | 20.66 | 68.87 | 6.19 | 0.85 | 0.08 | 11.13 | 7.92 | 25.28 |
| **Ina** | 8.04 | 17.99 | 73.10 | 5.24 | 0.59 | 0.08 | 7.65 | 10.63 | 17.6 |
| **Kandi** | 7.57 | 12.25 | 79.37 | 5.08 | 0.41 | 0.06 | 7.24 | 6.67 | 25.09 |
| **Natitingou** | 14.88 | 23.92 | 60.57 | 5.58 | 1.18 | 0.12 | 9.57 | 11.88 | 17.14 |
| **Parakou** | 9.35 | 17.64 | 72.41 | 5.90 | 0.81 | 0.08 | 9.76 | 9.17 | 19.85 |
| **Source**: Soil analyses at the Laboratory of Soil Science, Republic of Benin. | | | | | | | | | |

**Table S3. Agronomic traits of the eleven genotypes used in this study**

| **Genotypes** | **Origin** | **Germination rate (%)** | **Average plant height** | **Average days to 50% maturity** | **Grain yield (kg ha^-1^)** |
| --- | --- | --- | --- | --- | --- |
| **B12** | Benin | 55 | 70.52 | 104.50 | 263.78 |
| **B33** | Benin | 20 | 73.10 | 102.75 | 324.02 |
| **B37** | Benin | 40 | 72.32 | 89.50 | 313.37 |
| **BF1** | Burkina Faso | 70 | 78.63 | 92.50 | 300.99 |
| **G4** | Guinea | 60 | 89.13 | 101.50 | 227.71 |
| **G31** | Guinea | 60 | 84.83 | 103.75 | 237.38 |
| **G32** | Guinea | 60 | 85.08 | 101.50 | 217.52 |
| **M5** | Mali | 70 | 79.67 | 77.67 | 387.72 |
| **M14** | Mali | 65 | 73.22 | 78.33 | 478.15 |
| **M15** | Mali | 75 | 73.00 | 95.67 | 411.85 |
| **N35** | Niger | 95 | 77.11 | 85.33 | 417.78 |
| **Sources:** Own data and Ibrahim Bio Yerima et al. (2020). | | | | | |

**Table S4. Mean values of days to flowering (DTF), days to maturity (DTM) and grain yield (GRY) by environments studied.**

|  | **Days to flowering (DTF)** | **Days to maturity (DTM)** | **Grain yield (GRY)** |
| --- | --- | --- | --- |
| **Mean performance** | 84.5** | 116.02*** | 352.71*** |
| **Boukoumbe** | 87.4a | 117.37a | 106.56bc |
| **Gogounou** | 81.83ab | 104.6b | 794.52a |
| **Ina** | 86.39a | 118.23a | 715.54a |
| **Kandi** | 86.57a | 115.96a | 303.99b |
| **Natitingou** | 76.54b | 118.38a | 95.47bc |
| **Parakou** | 86.83a | 122.17a | 22.36c |

DTF in DAS, DTM in DAS, GRY in kg ha^-1^. Values (mean) within a column followed by no letters or the same letter are not significantly different according to the Tukey test at p ≤ 0.05.

**Table S5. Mean values, AMMI stability value (ASV) and ranking for the genotypes studied.**

|  | **Days to flowering (DTF)** | | | **Days to maturity (DTM)** | | | **Grain yield (GRY)** | | |
| --- | --- | --- | --- | --- | --- | --- | --- | --- | --- |
| **Genotype** | **Mean** | **ASV** | **Rank** | **Mean** | **ASV** | **Rank** | **Mean** | **ASV** | **Rank** |
| **B12** | 91.67 | 2.45 | 7 | 125.13 | 4.99 | 11 | 485.35 | 23.0 | 8 |
| **B33** | 90.46 | 4.73 | 11 | 124.36 | 1.35 | 2 | 329.807 | 27.5 | 9 |
| **B37** | 96.4 | 1.3 | 4 | 127.2 | 0.77 | 1 | 268.268 | 13.4 | 3 |
| **BF1** | 84.73 | 2.95 | 8 | 117.6 | 2.18 | 3 | 433.938 | 18.7 | 6 |
| **G31** | 92.35 | 1.89 | 5 | 121.29 | 4.1 | 9 | 480.282 | 7.83 | 1 |
| **G32** | 90.71 | 2.09 | 6 | 118.29 | 3.08 | 5 | 460.012 | 40.6 | 11 |
| **G4** | 89.07 | 4.31 | 10 | 121.2 | 4.19 | 10 | 256.038 | 16.2 | 5 |
| **M14** | 72.77 | 3.57 | 9 | 103.35 | 3.76 | 8 | 262.35 | 8.59 | 2 |
| **M15** | 69.87 | 0.95 | 1 | 104 | 2.4 | 4 | 172.123 | 13.8 | 4 |
| **M5** | 73.77 | 1.17 | 3 | 104.59 | 3.67 | 7 | 257.235 | 21.9 | 7 |
| **N35** | 81.11 | 1.01 | 2 | 114 | 3.36 | 6 | 443.546 | 29.5 | 10 |
| **Mean:** Mean values, **ASV:** AMMI stability value, **Rank:** Ranking. | | | | | | | | | |

**Table S6. Selection differential of the weighted average of absolute scores (WAASB) index for days to flowering (DTF), days to maturity (DTM) and grain yield (GRY) in the fonio genotypes studied.**

| **Traits** | **Factors** | **Xo** | **Xs** | **SD** | **SD%** |
| --- | --- | --- | --- | --- | --- |
| **Days to flowering (DTF)** | FA 1 | 45.5 | 71.2 | 25.7 | 56.4 |
| **Days to maturity (DTM)** | FA 1 | 46.5 | 69.6 | 23.1 | 49.6 |
| **Grain yield (GRY)** | FA 1 | 59.4 | 38.9 | -20.6 | -34.6 |
| **Xo:** original mean, **Xs:** mean of the selected accessions, **SD:** selection differential. | | | | | |

**Table S7. Environmental loadings for the factor analytic model.**

| **Environments** | **Days to flowering (DTF)** | | **Days to maturity (DTM)** | | **Grain yield (GRY)** | |
| --- | --- | --- | --- | --- | --- | --- |
|  | **FA1** | **FA2** | **FA1** | **FA2** | **FA1** | **FA2** |
| **Boukoumbe** | **7.32** | 3.62 | **4.27** | 0.43 | **17.4** | 15.56 |
| **Gogounou** | **6.29** | -4.86 | **6.68** | 1.25 | **87.24** | -45.76 |
| **Ina** | **6.85** | 0.46 | **14.42** | 0.25 | **80.63** | 40.7 |
| **Kandi** | **10.02** | 0.58 | **10.72** | -3.83 | 17.04 | **24.95** |
| **Natitingou** | **10.55** | -2.37 | **5.7** | 3.2 | 1.42 | **17.41** |
| **Parakou** | **12.51** | 1.61 | **11.16** | 0.81 | -1.75 | **7.02** |
| **FA1:** Factor 1, **FA2:** Factor 2, The highest values of the environmental loadings of the factorial model are highlighted in bold. | | | | | | |
